# The heterochromatin landscape in migrating cells and the importance of H3K27me3 for migration-associated transcriptional changes

**DOI:** 10.1101/344887

**Authors:** Tamar Segal, Mali Salmon-Divon, Gabi Gerlitz

## Abstract

H3K9me3, H3K27me3 and H4K20me1 are epigenetic markers associated with chromatin condensation and transcriptional repression. Previously, we found that migration of melanoma cells is associated with and dependent on global chromatin condensation that includes a global increase in these markers. Taken together with more recent reports by others suggest it is a general signature of migrating cells. Here, to learn about the function of these markers in migrating cells we mapped them by ChIP-seq analysis. This analysis revealed that induction of migration leads to expansion of these markers along the genome and to an increased overlapping between them. Significantly, induction of migration led to a higher increase in H3K9me3 and H4K20me1 signals at repetitive elements than at protein-coding genes, while an opposite pattern was found for H3K27me3. Transcriptome analysis revealed that 182 altered genes following induction of migration, of which 33% are dependent on H3K27me3 for these changes. H3K27me3 was also required to prevent changes in the expression of 501 other genes upon induction of migration. Taken together our results suggest that heterochromatinization in migrating cells is global and not restricted to specific genomic loci and that H3K27me3 is a key component in executing a migration-specific transcriptional plan.

## Introduction

Cell migration is one of cancer hallmarks that is fundamental for metastasis formation (Hanahan and Weinberg 2011). Tumor cell migration is highly dependent on cytoplasmic changes in the cytoskeleton and in the activity of motor proteins (Etienne-Manneville 2013; Rottner and Stradal 2011; Ouderkirk and Krendel 2014), but more recently alterations in chromatin structures were also found to be required (Gerlitz and Bustin 2011). Chromatin basic repetitive packaging unit is the nucleosome, which is composed of 147bp of DNA that are wrapped around an octamer of histones (McGinty and Tan 2014). Nucleosomes are organized in higher order structures, of which relatively de-condensed and transcribed regions are termed euchromatin, while more condensed and non-transcribed regions are termed heterochromatin. Euchromatin and heterochromatin are decorated by different levels and/or types of epigenetic marks such as histone modifications. Heterochromatin-associated histone modifications include H3K9me3, H3K27me3 and H4K20me1 (Mozzetta et al. 2015). H3K9me3 is enriched in constitutive heterochromatin regions such as repetitive elements and pericentromeric regions, where it is important for their repression (Peters et al. 2001). Still, H3K9me3 was also found at promoters of repressed genes as well as at some active genes (Barski et al. 2007). H3K27me3 accumulates over cell-type specific repressed genes (facultative heterochromatin) and the inactivate X chromosome (Barski et al. 2007; Margueron and Reinberg 2011, 2; Rougeulle et al. 2004). H4K20me1 is associated with repression of the inactive X chromosome and specific genes, but it was also found to accumulate downstream from the TSS of highly active genes (Beck et al. 2012; Kohlmaier et al. 2004). The inconsistent observations regarding H4K20me1 may arise from various cross-talks of H4K20me1 with other epigenetic marks and the fact that H4K20me1 is an initial step in the process of generating the highly repressive mark H4K20me3.

Previously we found that migration of mouse melanoma cells is associated with and dependent on global chromatin condensation that includes more than a two-fold increase in the levels of H3K9me3, H3K27me3 and H4K20me1 (Gerlitz and Bustin 2010; Gerlitz et al. 2007; Maizels et al. 2017). More recently, migration-associated global chromatin condensation was reported in primary and transformed T-cells (Zhang et al. 2016b) and primary tenocytes (Zhang et al. 2016a). Reliance of cell migration on chromatin condensation has been reported in various types of cells including lung cancer cells (Chen et al. 2010), embryonic fibroblasts (Kottakis et al. 2011), breast adenocarcinoma cells (Fu et al. 2012; Kokura et al. 2010; Yokoyama et al. 2013), colorectal cancer cells (Yokoyama et al. 2013), prostate cancer cells (Yu et al. 2017), glioma cells (Spyropoulou et al. 2014), chondrosarcoma cells (Girard et al. 2014), epidermal cancer stem cells (Adhikary et al. 2015), primary tenocytes (Zhang et al. 2016a) and primary and transformed T-cells (Zhang et al. 2016b). Thus, the dependence of tumor cell migration on chromatin condensation is a ubiquitous mechanism. Still, the exact roles of heterochromatin formation in migrating cells are not fully understood.

Here, to gain a mechanistic insight on the roles of heterochromatin in migrating cells we mapped the changes in heterochromatin spread and in the transcriptome in melanoma cells in response to migration signals by ChIP-seq and RNA-seq, respectively. Induction of migration was associated with more diffused distribution of H3K9me3, H3K27me3 and H4K20me1 that generated lower number of peaks, but had a higher over-lapping between the different histone modifications. Following induction of migration, H3K9me3 and H4K20me1 accumulated to a higher degree in repetitive regions, while H3K27me3 re-distributed towards genes. In parallel, we identified 182 genes with altered RNA levels of which one third were dependent on migration-induced methylation of H3K27.

## Results

Induction of migration leads to a global increase of 2-4-fold in the levels of the heterochromatin markers H3K9me3, H3K27me3 and H4K20me1, as we previously detected by immunostaining (Maizels et al. 2017; Gerlitz and Bustin 2010; Gerlitz et al. 2007). To learn about the mechanistic role of these histone modifications in the migration process, we mapped their relative distribution along the genome by ChIP-seq. Migration of the B16-F1 mouse melanoma cells was induced by the wound healing assay (Liang et al. 2007), in which multiple scratches were performed in each plate to generate large numbers of migrating cells. This way of induction of migration enabled us to receive an enriched population of migrating cells as measured by a 2-5-fold increase in the levels of H3K9me3 and H3K27me3 at the promoters of *E-cadherin, Gapdh* and *Line* (Sup. Fig. 1).

H3K9me3, H3K27me3 and H4K20me1 ChIP-seq mapped reads were at the range of 27-42 million and the coverage values at the range of 0.68-1.52 (Sup. Fig. 2a). As expected from the nature of heterochromatin modifications, more than 50% of the signals of these modifications did not accumulate at defined and short loci to form sharp peaks (Fig. 1a). Moreover, following indication of migration, this phenomenon further increased, resulting in accumulation of only 7.45%, 9.62% and 29.64% of the reads of H3K9me3, H3K27me3 and H4K20me1, respectively, inside peaks (Fig. 1a). In agreement, upon induction of migration the intensities of the peaks were reduced by 14-17% (Fig. 1a), and the number of identified peaks was reduced by 30-40%, while the number of enriched-peaks was reduced by more than 90% (Sup. Fig 2b). Significantly, upon induction of migration the average peak length of H3K9me3 and H4K20me1 was increased by 34% and 20%, respectively, while the average peak length of H3K27me3 was reduced by 20% (Fig. 1a). Taken together, the above analyses indicate on more diffused signals of H3K9me3, H3K27me3 and H4K20me1 upon induction of migration. This pattern suggests a possible increase in the degree of overlap between the three modifications following induction of migration. Indeed, in migrating cells the correlation between these modifications increased significantly to 0.73 between H3K9me3 and H3K27me3, 0.77 between H3K9me3 and H4K20me1 and 0.69 between H3K27me3 and H4K20me1 (Fig. 1b). The increased correlation between the different modifications occurred over any evaluated genomic element (promoters, enhancers etc.) (Sup. Fig. 3).

**Figure 1:**
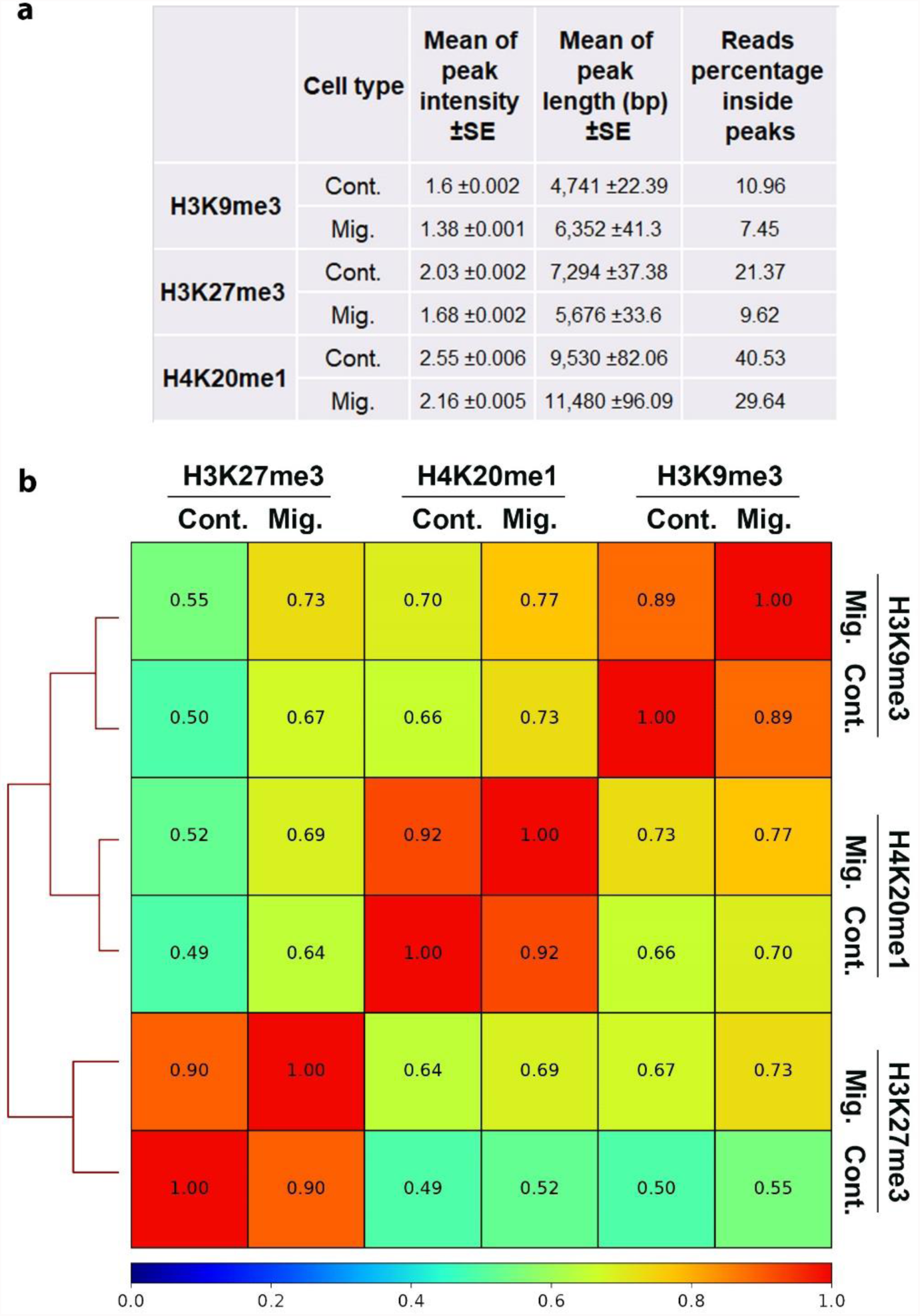
The patterns of the ChIP-seq signals of H3K9me3, H3K27me3 and H4K20me1 upon induction of migration. (**a**) Mean ± SE of ChIP-seq peak intensities and lengths of H3K9me3, H3K27me3 and H4K20me1 in control (Cont.) and migrating (Mig.) cells. Reads percentage inside peaks are the percentage of ChIP-seq mapped reads of the indicated heterochromatin markers that are localized inside peaks. Statistical significance was calculated between control cells to migrating cells by Wilcoxon rank sum test, p.val<2.2*10-16. **(b)** Genome wide correlation between the three heterochromatin markers. Spearman correlation coefficients of the ChIP-seq signals were calculated from reads coverage of consecutively equally sized 10kb bins genome-wide.

To assess which genomic regions are more prone to be affected in migrating cells by each of the above modifications, we counted the number of mapped reads overlapping specific genomic regions and calculated them as percentage of the total mapped reads. The functional genomic regions in our analysis included protein coding genes, non-coding RNA, enhancers, promoters and repetitive elements. The relative distribution of H3K9me3 and H4K20me1 mapped reads increased at repetitive elements by 2% and 19%, respectively, while decreased at protein coding genes by 1% and 11%, respectively. On contrary, the amount of H3K27me3 mapped reads increased by 7% at protein coding genes, while decreased by 3% at repetitive elements (Fig. 2a). This pattern became more definite when the relative distribution of differential peaks that fall inside different genomic elements was calculated: The relative distribution of nucleotides in differential-peaks of H3K9me3 and H4K20me1 at repetitive elements increased by 77% and 546%, respectively, while decreased at protein coding genes by 23% and 37%, respectively. On contrary, the nucleotides in differential-peaks of H3K27me3 increased by 92% at protein coding genes, while decreased by 54% at repetitive elements (Fig. 2b). To verify these results, we analyzed the average signal distribution of these modifications across different types of repetitive elements and across protein coding genes. In agreement with the previous analysis, the signals of H3K9me3 and H4K20me1 were higher across LINE, SINE, LTR and DNA transposons in migrating cells than in control cells, while the signal of H3K27me3 was lower across the same repetitive elements in migrating cells than in control cells. An opposite pattern emerged over protein coding genes: reduced levels of H4K20me1 together with increasing levels of H3K27me3 in migrating cells than in control cells (Fig. 3a).

**Figure 2:**
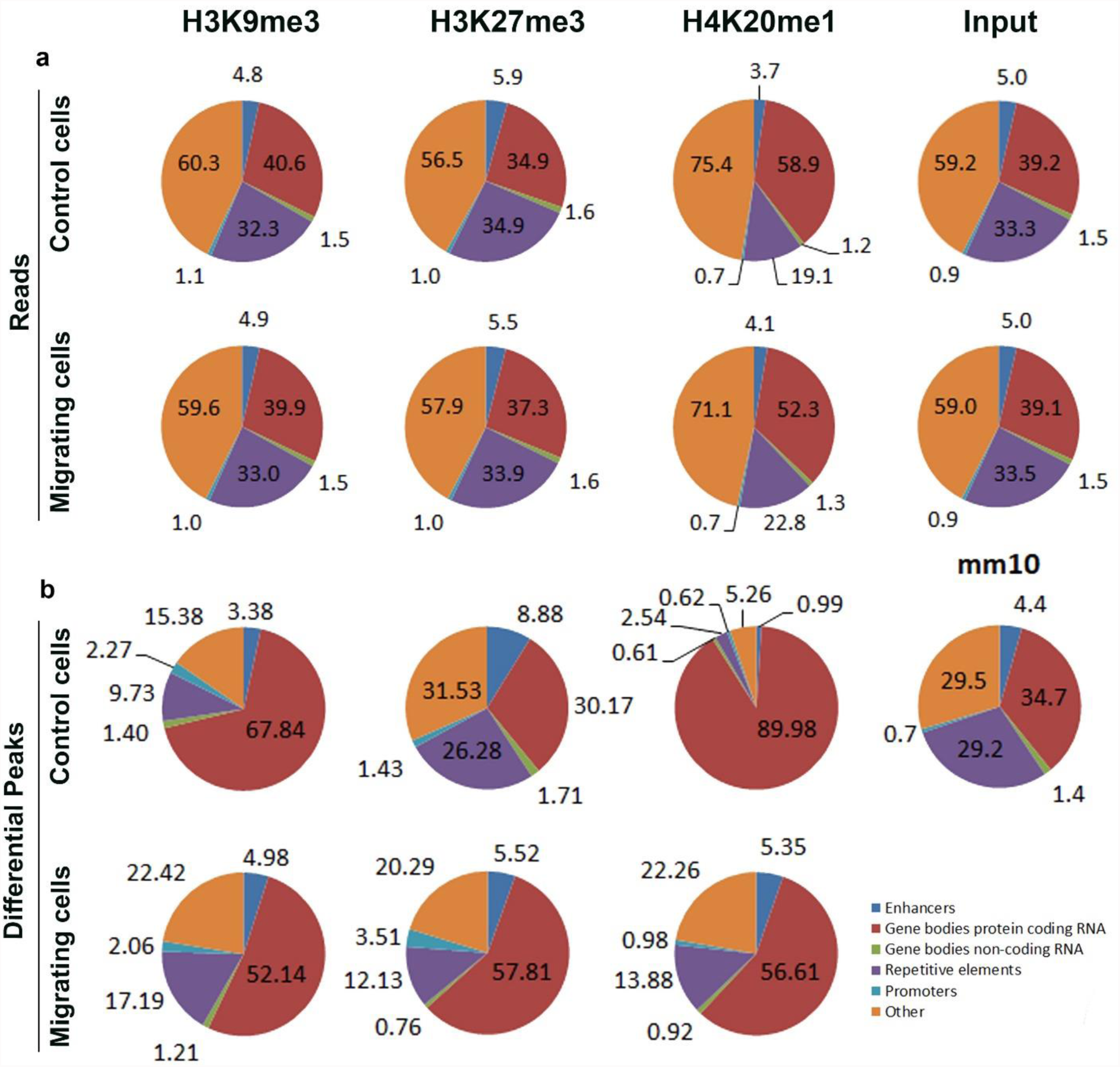
Relative distribution of ChIP-seq signal across various genomic elements in control and migrating cells. The relative distribution of ChIP-seq reads (a) and ChIP-seq differential peaks (b) across the indicated genomic elements was calculated for the input and the three heterochromatin markers in control cells and in migrating cells.

**Figure 3:**
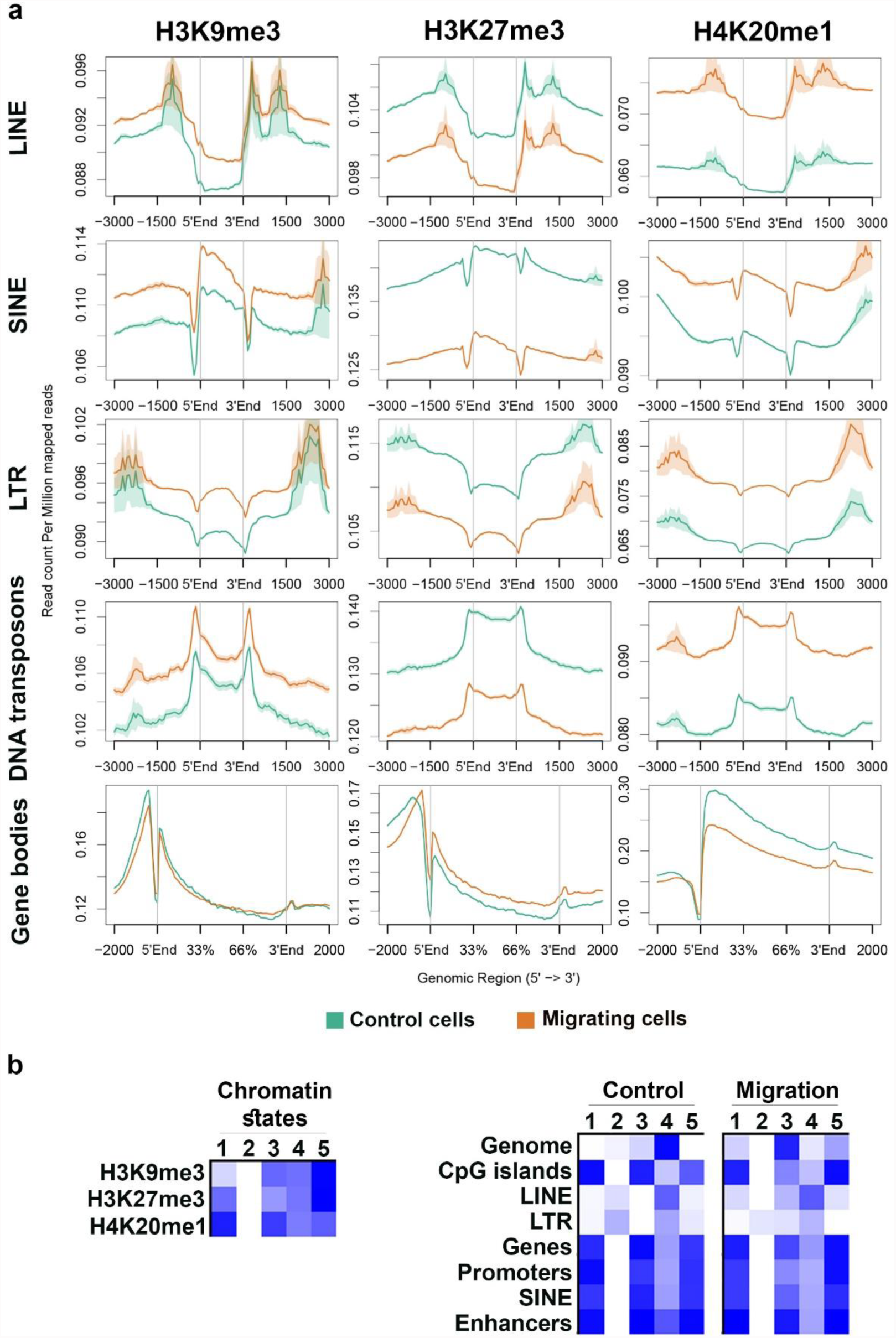
Distinct and combinatorial pattern of heterochromatin markers over genomic elements. (**a**) Distribution of H3K9me3, H3K27me3 and H4K20me1 ChIP-seq signals in control cells (green line) and in migrating cells (orange line) across gene bodies and the repetitive elements LINE (n=671156), SINE (n=682126), LTR (n=646284) and DNA transposons (n=79756). In each graph, the region between 5’End and 3’End represents the analyzed element. (**b**) Five combinatorial chromatin states were defined using the three heterochromatin markers. The left panel describes the chromatin states as represented by the emission coefficients in ChromHMM model, the right panel describes the relative enrichment of states coverage across the whole genome and over different genomic regions, in control and migrating cells.

Classifying the combinatorial pattern of the above modifications using a Hidden-Markov-Model (HMM) based approach revealed that upon migration there is a reduction in the genomic coverage by regions with similar levels of the three markers (state no. 4, Fig. 3b), while there is an increase in the percentage of genomic regions that are decorated by all three markers, but at differential levels (states no. 1, 3 and 5, Fig. 3b). Interestingly, regions free of all three markers occur in repetitive elements (LINE and LTR), but their percentage decreases upon induction of migration (state no. 2, Fig. 3b). Importantly, upon induction of migration the changes in the combinations of the modifications across promoters and gene bodies were much smaller than the changes over the whole genome, thus raising the question if there are any transcriptional changes in migrating cells.

To answer this question, we determined the changes in the transcriptome upon induction of migration using RNA-seq (Sup. Fig. 4). Differential expression analyses identified 801 genes with altered expression levels (FDR<0.05) in cells that were induced to migrate for 3h: 465 genes were up-regulated and 336 genes were down-regulated. Out of them, 182 genes were altered with a fold change of >1.3: 129 up-regulated genes and 53 down-regulated genes (Sup. Table 1 and Sup. Fig. 5). Surprisingly, although induction of migration induces a global increase in chromatin condensation, higher number of genes were upregulated than down-regulated. To search for common functions of the altered genes, genes found to be differentially expressed with FDR >0.05 and fold change >1.3 were subjected to Ingenuity Pathway Analysis (IPA). As shown in Fig. 4, the most significantly enriched pathways included ones that are known to be associated with tumor cell proliferation and migration such as the TGF-β, IGF-1 and ERK5 signaling pathways (Gabellini et al. 2009, 8; Giampieri et al. 2009; Ning et al. 2011; Pang et al. 2016; Simões et al. 2016, 5) and pathways involved in energy generation: glycolysis and pyridoxal 5’-phosphate (PLP) salvage pathway. PLP is an active form of vitamin B6, which is important for the metabolism of carbohydrates, amino acids and fats, the generation of the methyl donor S-adenosylmethionine (SAM) and in neutralizing oxidative stress (Bird 2018; Anderson et al. 2012). Activation of oxidative stress response was indicated also by the up-regulation of NRF2 targets (Fig. 4). Analysis of upstream regulators and downstream affected functions identified a strong link to cell migration (Sup. Fig. 6).

**Figure 4:**
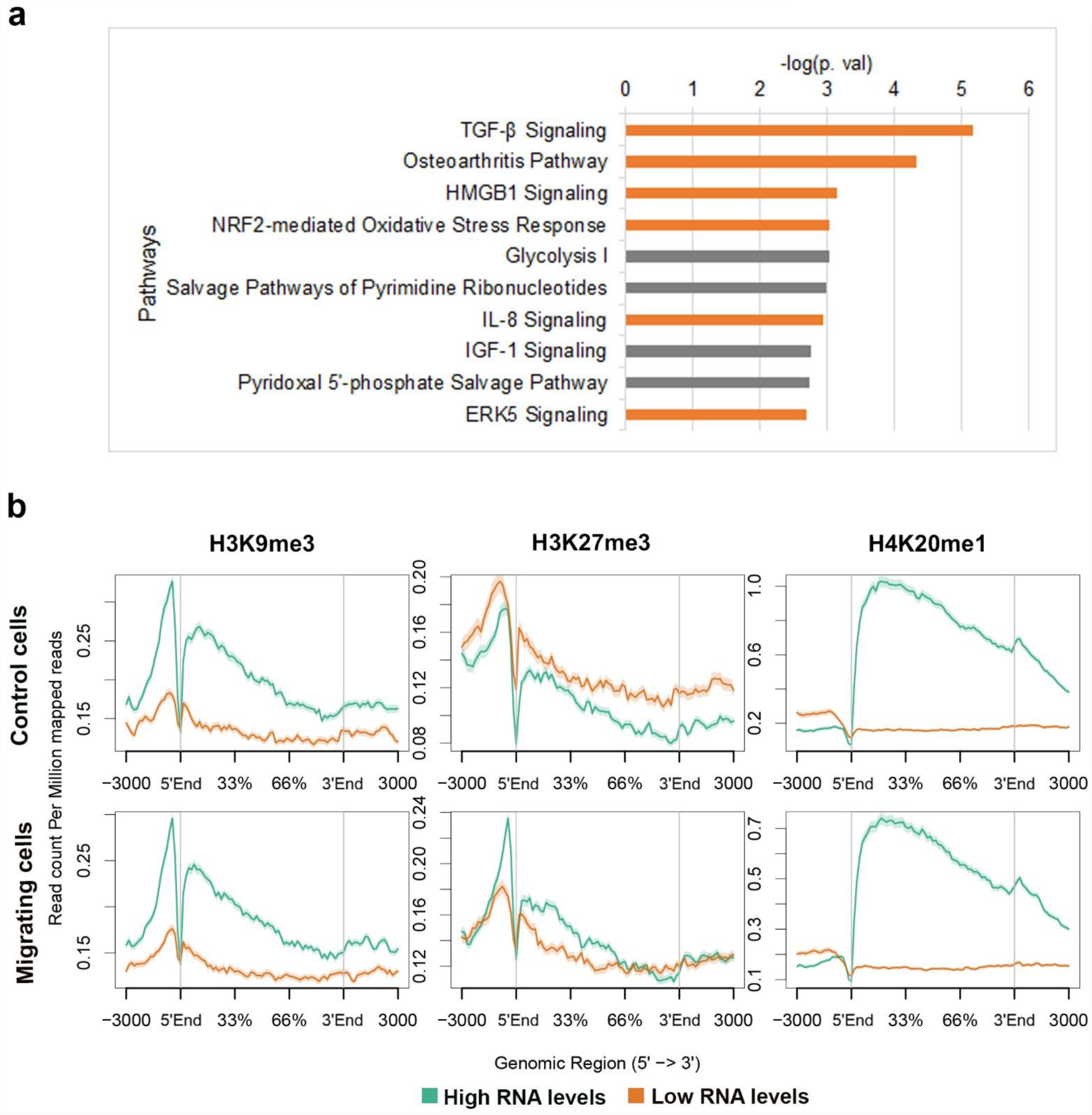
Altered gene expression upon induction of migration. (**a**) Top 10 pathways that are altered upon induction of migration as were identified by IPA using RNA-seq data collected from control cells and migrating cells (n=5). Only genes that were differentially expressed by factor of >1.3 and FDR<0.05 were included in the analysis. Orange bars represent up-regulated pathways and grey bars represent altered pathways in which the altered genes did not have a uniform trend. (**b**) Distribution of heterochromatin markers across highly (green) and lowly (orange) abundant genes. Genes with average expression level higher than 1 cpm were selected, out of these, the highest or lowest 500 expressed genes were chosen. Genes bodies are presented between the 5’End and the 3’End marks. Additional 3000bp upstream and downstream of the genes are plotted.

To evaluate the correlations between H3K9me3, H3K27me3 and H4K20me1 and transcription in our system, we first assessed the intensities of these modifications over genes that exhibit different expression levels. All three evaluated modifications did not accumulate over non-expressed genes (Sup. Fig. 7-9), suggesting a different repression mechanism for these genes, such as DNA methylation. In expressed genes, H4K20me1 and H3K9me3 patterns were similar between control and migrating cells: H4K20me1 levels across promoter regions correlated inversely with gene expression levels, whereas the contrary was observed across gene bodies (Fig. 4b and Sup. Fig. 9). Surprisingly, H3K9me3 signal intensity correlated positively with gene expression levels (Fig. 4b and Sup. Fig. 7). In contrast to H4K20me1 and H3K9me3, H3K27me3 association with gene expression levels overturned upon induction of migration: In control cells, as expected, H3K27me3 signal intensity correlated inversely with gene expression levels. However, in migrating cells, H3K27me3 correlated positively with gene expression levels (Fig. 4b, Sup. Fig. 8).

Out of the three analyzed histone modifications, only H3K27me3 displayed increased levels over genes upon induction of migration (Fig. 3) and its reverse correlation with gene expression levels was turned-over upon induction of migration (Fig. 4b and Sup. Fig. 8). Therefore, we analyzed which migration-associated transcriptome changes are dependent on H3K27me3 by RNA-seq of migrating cells that were treated with the Ezh2-specific inhibitor, GSK343, for 3 h. This analysis revealed that 33% of the migration-altered genes were not changed once GSK343 was added, indicating they are H3K27me3-dependent genes. The 67% of the migration altered genes that were changed also in migrating cells treated with GSK343 were termed H3K27me-independent genes (Fig. 5a, Sup. Table 2, Sup. Fig. 4 and Sup. Fig. 10). Significantly, most of the signaling pathways altered in migrating cells such as TGF-β and ERK5 were H3K27me-dependent, while the most significant H3K27me independent-pathways were glycolysis and NRF2-mediated response (Fig. 5b). In addition, 501 genes that normally do not change upon induction of migration did change once H3K27 methylation was inhibited (Fig. 5, Sup. Table 2 and Sup. Fig. 10). These genes are mainly involved cholesterol metabolism. Thus, H3K27 methylation in migrating cells is important not only for establishing a new transcriptional plan, but also to limit the transcriptional changes to specific genes. Analyzing the levels of H3K27me3 across the migration-altered genes revealed no obvious changes in H3K27me3 levels between control and migrating cells over both H3K27me-dependent and independent genes (Fig. 5c). Still, H3K27me3 levels at the promoters of the H3K27me-depedent genes were higher than those of the H3K27me-independent genes. On contrary, over the H3K27me-buffered genes a clear increase in H3K27me3 levels upon induction of migration was observed. Suggesting that the effect of H3K27me3 on the H3K27me-depednet genes may be indirect.

**Figure 5:**
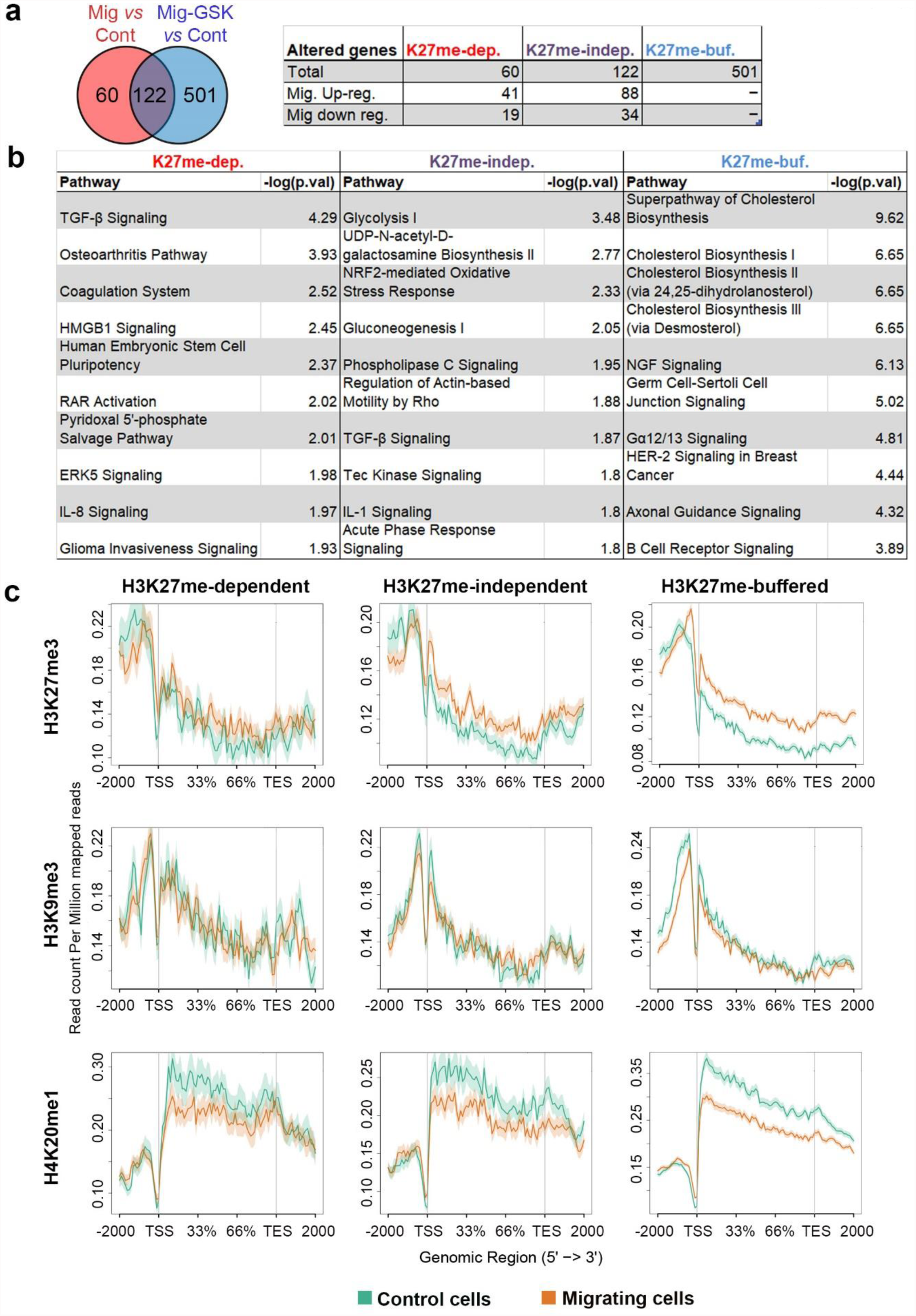
The dependence of migration-induced transcriptional changes on H3K27 methylation. (**a**) Altered genes upon induction of migration (fold change >1.3, FDR<0.05) that were detected in untreated migrating cells and in migrating cells treated with GSK343 are represented by a Venn diagram. Overlapping genes are termed “H3K27me-independent”, genes altered only when migrating cells were compared to control cells are termed “H3K27me-dependent” and differentially expressed genes only in GSK343-treated migrating cells are termed “H3K27me-buffered”. (**b**) Top 10 pathways altered according to the “H3K27me-independent” gene list, the “H3K27me-dependent” gene list and the “H3K27me-buffered” gene list as identified by IPA. (**c**) Distribution of H3K27me3, H3K9me3 and H4K20me1 signals across “H3K27me-dependent”, “H3K27me-independent” and “H3K27me-buffered” genes in control cells (green) and in migrating cells (orange). The region between 5’End and the 3’End represent genes bodies. Additional 2000bp upstream and downstream of the genes are plotted.

Finding that 67% of the migration-altered transcripts are independent of H3K27 methylation means that additional mechanisms are involved in their regulation. To learn if alterations in their RNA stability levels are involved, we determined the transcriptome of control and migrating cells following 3 h of RNA polymerase II inhibition by DRB. Calculation of the ratio between the mRNA levels in DRB-treated migrating cells and DRB-treated control cells identified 653 downregulated genes and 365 upregulated genes having decreased and increased RNA stability upon induction of migration, respectively (Fig. 6a, Sup. Table 3, Sup. Fig. 4 and Sup. Fig. 11). Notably, whereas only 10% of the H3K27me-dependent genes had migration-dependent changes in their RNA stability levels, 23% of the H3K27me-independent genes had migration-dependent changes in their RNA stability levels. IPA analysis revealed that genes with increased RNA stability in migrating cells are mainly involved in translational control and energy production, while genes with decreased RNA stability in migrating cells are mainly involved in cell cycle control and DNA damage response (Fig. 6b). Suggesting that regulation at the levels of RNA stability and translation rate may be highly important for cell migration.

**Figure 6:**
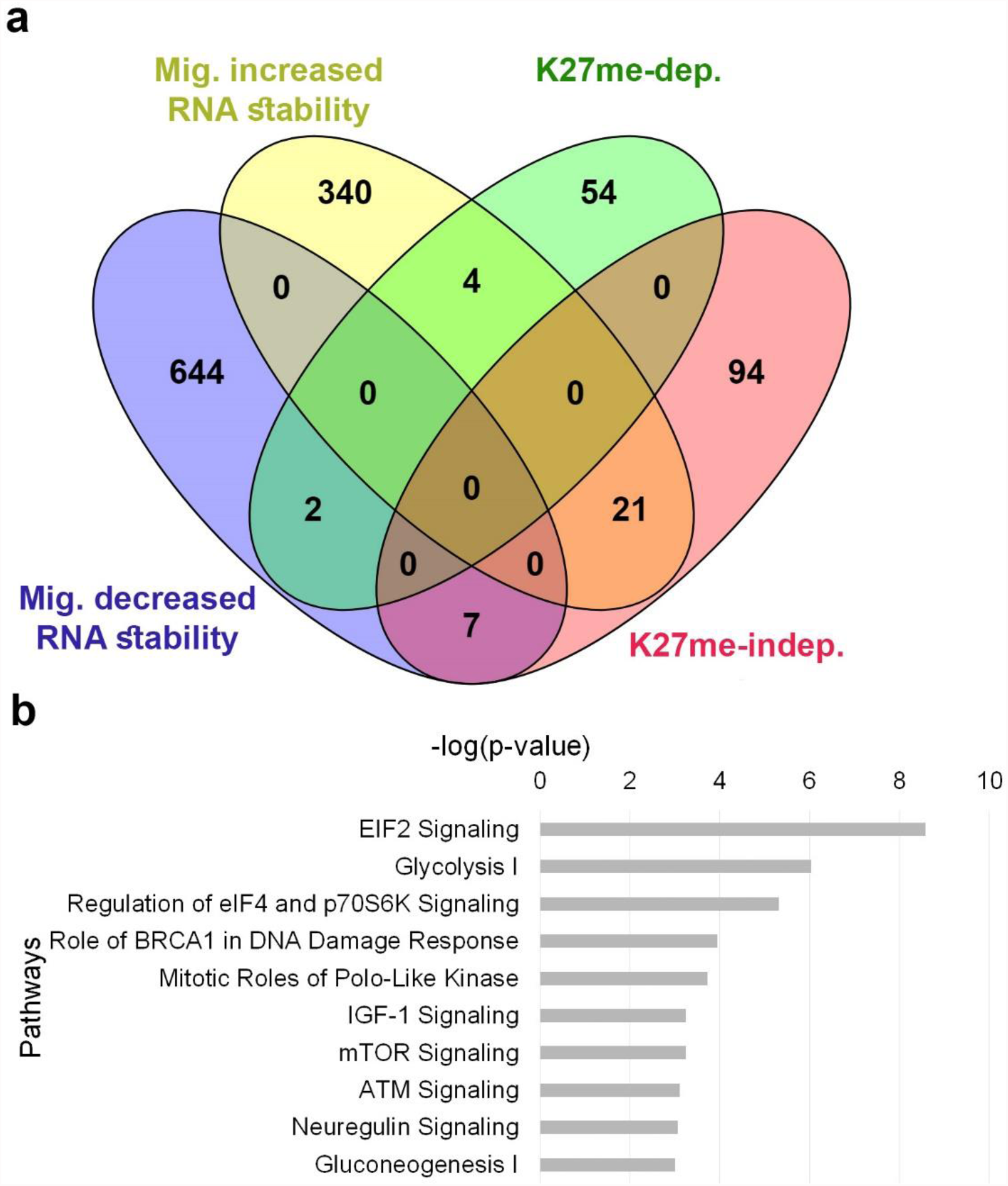
Migration-induced changes in RNA stability. Altered genes upon induction of migration were detected in the presence of DRB, an RNA polymerase II inhibitor (fold change >1.3, FDR < 0.05). Increased RNA stability is detected by upregulation of genes in DRB-treated migrating cells compared to DRB-treated control cells, and decreased RNA stability by downregulation. (**a**) A Venn diagram representing the overlap between genes with increased or decreased RNA stability levels upon induction of migration to H3K27me-dependent and independent genes. (**b**) Top 10 pathways of the genes with altered RNA stability levels upon induction of migration as identified by IPA.

## Discussion

In this study we found that upon induction of migration, the increase in heterochromatin markers does not occur in narrow regions to generate peaks, but rather spread over larger genomic regions. Migration signals alter the RNA levels of 182 genes that are involved in signaling pathways such as TGFβ and ERK5, but also in glucose metabolism and oxidative stress response. One third of the transcriptome alterations are dependent on methylation of H3K27, while altered genes in H3K27me-indepednent manner are more prone to regulation at their RNA stability level.

ChIP-seq protocols depend on the assumption that the overall yields of DNA are identical per cell under different conditions. Accordingly, the same amount of total DNA is taken for analysis, and the resulting data are normalized to each other so that the total amounts of signals are identical. Since induction of migration is associated with a dramatic increase in the amount of heterochromatin, ChIP doesn’t yield the same amount of DNA in control and migrating cells. Hence, the DNA normalization done during library preparation reduces the quantitative difference between the two conditions, which may explain, the lack of global increase in heterochromatin markers upon induction of migration in our ChIP-seq experiments. Therefore, our analysis has been mainly focused on the changes in signal distribution across genomic features upon induction of migration.

Induction of migration led to a more diffuse distribution of H3K9me3, H3K27me3 and H4K20me1 along the genome as measured by reduced number of ChIP-seq peaks and ChIP-seq signals within peaks (Fig. 1 and Sup. Fig. 2). In agreement with this pattern, the over-lapping between the different markers increased dramatically (Fig. 1). These changes were accompanied by a shift in the distribution of these markers between the various genomic elements: H3K9me3 and H4K20me1 signals shifted from protein-coding regions to repetitive elements while the H3K27me3 signal shifted in the opposite direction. These shifts were seen in whole reads analysis and become much more dramatic when analyzing the reads that fell only within differential peaks (Fig. 2). These patterns suggest that although the overall signals of each of these markers is increased by 2-4 fold as previously measured by immunostaining (Gerlitz and Bustin 2010; Gerlitz et al. 2007; Maizels et al. 2017) the effects on transcription may be moderate since the signals do not accumulate at narrow regions. Indeed, transcriptome analysis revealed that upon induction of migration only 182 genes were altered with a fold change of >1.3, while only 30% of them were down regulated.

Reports on the transcriptome of migrating cells are quite scarce (Dayem et al. 2003; Demuth et al. 2008; Fitsialos et al. 2007). In previous reports the transcriptome was evaluated by microarrays that had a limited genomic coverage. Analyzing our RNA-seq data by IPA identified the activation of TGFβ, IL-8 and ERK5 signaling pathways that were found to be activated in previous transcriptome analyses of migrating cells (Dayem et al. 2003; Demuth et al. 2008; Fitsialos et al. 2007) (Fig. 4). These pathways are well established regulators of cancer cell migration and invasion (Gabellini et al. 2009; Giampieri et al. 2009; Ning et al. 2011; Pang et al. 2016; Simões et al. 2016). In addition, we found changes in glucose metabolism that includes up-regulation of three glycolytic enzymes (Glucose-6-Phosphate Isomerase, Phosphofructokinase and Aldolase C) that may be required for increased energy production in migrating cells. The increase in oxidative stress response genes may be required due to the increase in ATP production.

In the literature, there are mixed reports regarding the association between H4K20me1 signal and gene expression levels (Beck et al. 2012; Kohlmaier et al. 2004). Our integrated analysis of ChIP-seq and RNA-seq data identified a positive-correlation between H4K20me1 levels downstream the TSS and gene expression levels, and negative-correlation between H4K20me1 signal intensity upstream the TSS and gene expression levels (Fig. 4 and Sup. Fig. 9). Thus, H4K20me1 may support transcription when localized downstream the TSS whereas it may be involved in transcriptional repression when localized upstream of the TSS.

Using an Ezh2 specific inhibitor we were able to find that methylation of H3K27 is required for the transcriptional regulation of 33% of the migration-altered genes as well as to prevent changes in 501 additional genes upon induction of migration (Fig. 5). Migration signals lead to activation of several transcriptional regulators such as SMAD4 and SP1 (Sup. Fig. 6) that have the potential to affect the transcription of several hundreds of genes. We hypothesize that the changes in the transcription of many of these genes are not required or may even interfere with the migration process, and H3K27me3 serves to prevent the binding or the activity of transcription factors at these genes.

An additional mechanism to prevent changes in the transcriptome beyond the 182 migration-altered genes is by changes in RNA stability levels. Our analysis identified 1018 genes with altered RNA stability level upon induction of migration (Fig. 6). However, the net mRNA level of most of these genes do not change in migrating cells, indicating on inverse changes in their transcription rate.

In search for some insight regarding the mechanism that controls the migration-induced changes in the H3K27me-independent genes we found that these genes are more susceptible to alterations in their RNA stability levels following induction of migration (Fig. 6).

In analyzing the importance of transcription for cell migration we found previously that active transcription is dispensable for migration during a time period of 3 h, whereas it is important for migration during longer period of time (11 h) (Gerlitz and Bustin 2010). Thus, the changes we identified here are the initial changes that will have an accumulative effect as the migration time increases. Still, heterochromatin is important for migration during a time period of 3 h as well (Gerlitz and Bustin 2010; Gerlitz et al. 2007). This may be due to the effect of chromatin condensation on the physical properties of the nucleus (Gerlitz and Bustin 2011, 2010; Bustin and Misteli 2016). Increased heterochromatin levels were shown to increase nuclear rigidity (Zhang et al. 2016a; Furusawa et al. 2015; Stephens et al. 2018), which may be important to protect the genome from mechanical damage that can occur during migration through confined spaces (Denais et al. 2016; Raab et al. 2016). Increased nuclear rigidity may also serve as a better anchoring point for the actin cytoskeleton that was suggested to use the nucleus to generate force during cell migration (Graham et al. 2018; Petrie et al. 2017). In support of the mechanical roles of migration-induced chromatin condensation are the increased spread of the heterochromatin markers over larger genomic regions. Thus our results support a model in which migration-induced chromatin condensation serves to alter both the transactional plan as well as the physical properties of the nucleus.

## Materials and Methods

### Cell culture

Mouse melanoma B16-F1 cells were grown in DMEM (D5796, Sigma-Aldrich, St. Louis MO, USA) supplemented with 10% FCS (04-007-1A Biological Industries, Beit Haemek, Israel), 0.292 mg/mL L-glutamine (03-020-1B, 1A Biological Industries, Beit Haemek, Israel) and 40 units/mL Penicillin-Streptomycin (03-031-1B, Biological Industries, Beit Haemek, Israel) at 37 OC, 7% CO_2_. For migration assays the cells plated on fibronectin-coated plates were grown to confluence. To induce migration, the cells were scratched at multiple sites, washed once with DMEM and incubated in growth medium at 37 OC and 7% CO_2_ for 3 h. Control cells were kept in similar conditions without being scratched. To inhibit the generation of H3K27me3 or to inhibit RNA polymerase II the cells were incubated in growth medium supplemented with 3 µM of GSK343 (SML0766, Sigma-Aldrich, Rehovot, Israel) or 0.1 mM DRB (D1916, Sigma-Aldrich, Rehovot, Israel). The inhibitors were added 3h before lysing the cells.

### ChIP-seq

Chromatin immunoprecipitation (ChIP) was performed as described previously (Aparicio et al. 2005) with the following modifications. Following cross linking with 1% PFA, cells were collected and aliquoted to 10^7^ cells. Nuclei were isolated by lysis buffer and DNA was collected following, fragmentation by 0.7 mU/μl of MNase (N3755, Sigma-Aldrich, Rehovot, Israel) at 37°c for 15 min and briefly sonication in RIPA buffer. IP was done using magnetic protein A/G beads (BioVision, 6527-1, CA, USA) coupled to 24µl of rabbit α H3K9me3 (Millipore, 07-442, CA, USA), 12µl of rabbit α H3K27me3 (Millipore, 07-449, CA, USA) or 24µl of rabbit α H4K20me1 (Millipore, 17-610, CA, USA). DNA was purified using QIAquick Gel Extraction kit (QIAGEN, 28704, Hilden, Germany) according to the manufacturer protocol and sequenced at the Technion Genome Center by Illumina HiSeq 2500 machine.

### RNA purification and RNA-seq

Total RNA was purified by the NucleoZOL kit (MACHEREY-NAGEL, 740404.200, Duren, Germany) according to the manufacturer’s instructions. Purified RNA was sent for poly A containing mRNA selection, library preparation and sequencing at the Technion Genome Center. Replicates number was five for untreated control and migrating cells and three for cells treated with GSK343 or DRB.

### Peak calling and peak analysis

Quality control checks of the raw sequence data were carried out using the FastQC tool ([CSL STYLE ERROR: reference with no printed form.]). Then, the Trim_galore ([CSL STYLE ERROR: reference with no printed form.]) tool that is based on cutadapt (Martin 2011) was used for adapters trimming and for removing of low quality bases from the ends of reads. The first four nucleotide which were of bad quality were trimmed from reads by fastX trimmer ([CSL STYLE ERROR: reference with no printed form.]). Cleaned, high quality reads were aligned to the mouse genome (build mm10) using Bowtie2 (Langmead and Salzberg 2012) with default parameters. PCR bias of duplicated reads were removed using “MarkDuplicates” command implemented by picard tool ([CSL STYLE ERROR: reference with no printed form.]). Broad peaks enriched in immunoprecipitation over input were identified by SICER (Xu et al. 2014) with “fragment size” of 350, “effective genome fraction” of 0.8 and “FDR” 0.05. SICER-df was applied to determine differential peaks enriched in migrating cells compared to control cells and vice versa. Operations on genomic intervals were performed using BEDTools (Quinlan and Hall 2010). The ChIP-seq data reported in this paper were deposited at Gene Expression Omnibus database with accession number SUB3771416.

### Calculation of ChIP-seq coverage and signal distribution across specific genomic location

Genomic location of Refseq genes and repetitive elements were downloaded from UCSC table browser (Tyner et al. 2017). Enhancers location were downloaded from HOMER (Heinz et al. 2010) and the genomic coordinates were converted to mm10 assembly using UCSC liftOver tool. Promoters were defined as 1000bp upstream of transcription start sites (TSS). Within each list, overlapping intervals were merged into a single feature that spans all of the combined features. In order to avoid intervals overlapping more than a single defined annotated element, we removed from each list nucleotides that belong to another annotation list by assigning the following priority: coding genes, non-coding genes, promoters, enhancers and repetitive elements. The coverage of ChIP signal across these regions was presented as percentage of reads mapping within genomic regions out of the total mapped reads without redundancy, or as percentage of bp mapping within genomic regions out of the total number of bp included within peaks or differential peaks, and was calculated using coverageBed (Quinlan and Hall 2010) and intersectBed (Quinlan and Hall 2010), respectively. Distribution of ChIP signal across functional genomic regions was done using ngs.plot (Shen et al. 2014).

### Correlation analysis and combinatorial pattern of heterochromatin markers

The entire genome was divided into non-overlapping equally sized bins (10kb), and the average ChIP score was calculated for each bin using multiBamSummary command from deepTools(Ramírez et al. 2016). Spearman correlation coefficients were computed using deepTool’s multiBamSummary and plotCorrelation commands. Hidden Markov model-based chromatin state definition was performed using the ChromHMM v1.14 (Ernst and Kellis 2017). Five state were used to segment the genome, and the enrichment of these states across different genomic feature was calculated.

### RNA-seq Analysis

Reads were aligned to the mouse genome (mm10) using TopHat (Trapnell et al. 2009) after removal of adapter sequences and critical examination of quality controls using Trim_galore ([CSL STYLE ERROR: reference with no printed form.]) with default parameters. The number of reads mapping each mouse gene (as annotated in Ensembl release GRCm38.86) was counted using the ‘intersection-nonempty’ mode of HTseq-count script (Anders et al. 2015). Differential expression analysis was performed using edgeR (Robinson et al. 2010) and Limma (Ritchie et al. 2015) packages from the Bioconductor framework (Reimers and Carey 2006). Briefly, features with less than 1 read per million in 3 samples were removed. The remaining gene counts were normalized using the TMM method, followed by voom transformation (Law et al. 2014). Linear models were used to remove batch effect and to find differentially expressed genes. Up and down regulated genes having FDR < 0.05 and fold change ≥ 1.3 were analyzed for pathway enrichment using the Ingenuity Pathway Analysis (Yu et al. 2016) (IPA). H3K27me3 dependency was detected by comparing genes found to be differentially expressed between migrating and control cells, to genes found to be differentially expressed between migrating cells treated with GSK343 to control cells. H3K27me3-dependent genes, are genes whose expression changes abolished upon treatment with GSK343, while H3K27me3-independent genes are those whose change in expression preserved even after treatment with GSK343.

### Distribution of heterochromatin signals across high and low abundant genes

Average gene expression was calculated across five replicates of RNA-seq samples for control cells and migrating cells and genes with average expression lower than 1 cpm (counts per million) reads were excluded. Genes were ranked based on their expression level, and the top and bottom 500 genes were chosen. The distribution of histone modifications signal across these genes was plotted using ngs.plot tool (Shen et al. 2014).

## Acknowledgments

We thank the Technion Genome Service and the Israel National Center for Personalized Medicine for their NGS services. Our research was supported by the Israel Cancer Research Fund (14-109-RCDA), the Israel Cancer Association (20150910) and Ariel Center for Applied Cancer Research.

## Author contributions

T.S., M.S.D. and G.G. planned the experiments, analyzed the data and wrote the manuscript. T.S. and G.G. performed the experiments.

